# The first complete mitochondrial genome of marigold pest thrips, *Neohydatothrips samayunkur* (Sericothripinae) and comparative analysis

**DOI:** 10.1101/361402

**Authors:** Vikas Kumar, Kaomud Tyagi, Shantanu Kundu, Rajasree Chakraborty, Devkant Singha, Kailash Chandra

## Abstract

The complete mitogenomes in order Thysanoptera is limited to subfamily Thripinae heretofore. In the present study, we sequenced the first mitochondrial genome of *Neohydatothrips samayunkur* (15,295 bp), a member of subfamily Sericothripinae. The genome was characterized by 13 protein-coding genes (PCGs), 22 transfer RNA genes (tRNAs), two ribosomal RNA genes (rRNAs) and three control regions (CRs). This mitogenome had two overlapping regions of 4 bp and twenty four intergenic spacers accounting for 165 bp. All the tRNA had typical cloverleaf secondary structures, except for *trnV and trnS* which lacked DHU stem and loop. The mitogenomes of *N. samayunkur* was highly rearranged with many unique features as compared to other thrips mitogenomes, *atp6* and *nad1* were terminated with TAG and TGA stop codons respectively; location of *trnL2*, *trnA*, *trnC*, and *trnV* was rearranged; and the first control region (CR1) was upstream of *nad6.* The phylogenetic analysis of 13 PCGs implementing maximum likelihood and Bayesian inference showed the clustering of *N. samayunkur* with *Scirtothrips dorsalis* supporting the *Scirtothrips* genus-group and Sericothripinae morphology based relationships. Generation of more mitogenomes from different hierarchical level in the order Thysanoptera is required to understand the gene rearrangements, phylogeny and evolutionary relationships.

## Introduction

The order Thysanoptera encompasses nine families in two recognized suborders, the Terebrantia and Tubulifera. The family Thripidae is the largest in Terebrantia and further subdivided in four subfamilies; Dendrothripinae, Panchaetothripinae, Sericorthripinae, and Thripinae^1^. The members of Sericothripinae have worldwide distribution and are usually associated with flowers^2,3^. This subfamily currently includes 168 species under three genera, *Neohydatothrips*, *Hydatothrips*, and *Sericothrips.* The marigold thrips, *Neohydatothrips samayunkur* is a pest of marigold (*Tagetes* spp.) with a worldwide distribution^4,5,6^. Recently, *N. samayunkur* is also suspected as vector for tospoviruses^5,7^. Integration of molecular data with morphology is required for fast and accurate species identification and phylogenetic relationship^8^. The efficacy of mitochondrial genes (cox1 and *16S rRNA)* have been found to be useful in identification of thrips species and their phylogenetic relationships^8,9^. However, the phylogenetic relationships below family level in thrips is still unclear and requires more molecular data^8,9,10^.

Mitochondrial (mt) genomes have been widely studied for phylogenetic and evolutionary studies in insects^11,12,13,14^. Insects have a typical circular mitochondrial genome, 14-19 kb in size, with 37 genes, including 13 protein-coding genes (PCGs), large and small ribosomal RNA (rRNAs), 22 transfer RNA (tRNAs) and variable number of A+T rich non-coding region. Till date, six highly rearranged mitogenomes of five thrips species (*Anaphothrips obscurus*, *Frankliniella intonsa*, *Frankliniella occidentalis*, *Scirtothrips dorsalis* and *Thrips imaginis*) are available in the GenBank database^15,16,17,18,19^. Nevertheless, the availability of mitogenomes in thrips is limited only to subfamily Thripinae so far. In this study, we sequenced the first complete mitochondrial genome of the marigold thrips, *N. samayunkur* under the subfamily Sericothripinae using next-generation sequencing (NGS) technology and comparative analysed with other thrips mitogenomes. We analysed the gene rearrangements, nucleotide composition, Relative Synonymous Codon Usage (RSCU), overlapping and intergenic spacer regions, and transitions and transversions saturation analysis of PCGs etc. Further, to infer the phylogenetic relationships, 13 PCGs of *N. samayunkur* and other published thrips mitogenomes were analysed using maximum likelihood (ML) and Bayesian inference (BI).

## Results and discussion

### Genome structure, organization and composition

The complete sequence of the mitochondrial genome of *N. samayunkur* (accession number MF991901) was 15,295 base pair (bp) in length. This mitogenome was greater than the genomes of *A. obscurus* (14,890 bp), *F. intonsa* (15,215 bp), *F. occidentalis* (14,889 bp) and *S. dorsalis* South Asia strain (SA1) (14,283 bp), but smaller than the genomes of *T. imaginis* (15,407 bp) and *S. dorsalis* East Asia strain (EA1) (15,343 bp) (Table S1). Like in other thrips species, the mitochondrial genome of *N. samayunkur* was represented by 37 genes, including 13 PCGs (*atp6*, *atp8*, *cox1*, *cox2*, *cox3*, *cob*, *nad1*, *nad2*, *nad3*, *nad4*, *nad4L*, *nad5*, and *nad6*), large and small ribosomal RNA (*rrnL* and *rrnS*) genes, 22 transfer RNA (tRNA) genes and three A+T-rich control regions (CRs) (Fig. 1). Thirty genes were observed on heavy (H) strand and seven genes on the light (L) strand (Table 1). The nucleotide composition revealed 77.42 % AT content (40.25% A + 37.17 % T) and 22.58% GC content (11.60 % C + 10.98% G) (Table 2). The mitogenomes of *N. samayunkur* was AT-rich as depicted in other thrips mitogenomes. In *N. samayuknur*, the AT compositions was observed to be highest 80.82% in tRNAs genes (1397 bp) followed by rRNAs genes (79.31%, 1808 bp), PCGs (77.13%, 10956 bp), and control regions (71.12%, 969 bp). Further, the mitogenome of *N. samayunkur* showed positive AT skewness (0.04) and negative GC skewness (-0.03). The AT skewness in other thrips mitogenomes sequenced so far, ranged from 0.15 (*T. imaginis)* to -0.02 (*A. obscurus*), while the GC-skewness from 0.01 (F*. occidentalis*) to -0.12 (*S. dorsalis* SA1). Moreover the positive AT skewness indicated the occurrence of more As than Ts that has also been reported in other four thrips species mitogenomes sequence except *A. obscurus* (-0.02). These skewness patterns of *N. samayunkur*, were also analysed in rRNAs genes, tRNAs genes, control regions and were found to be similar to the other thrips mitogenomes (Table 2). Further, sequence similarity searches among different thrips species revealed that *N. samayunkur* had highest similarity with *S. dorsalis* SA1 (75%) followed by *F. occidentalis* (74%), *F. intonsa* (73%), *T. imaginis* (71%), *S. dorsalis* EA1 (70%) and *A. obscurus* (70%) indicating a low level of homology.

**Figure 1.**
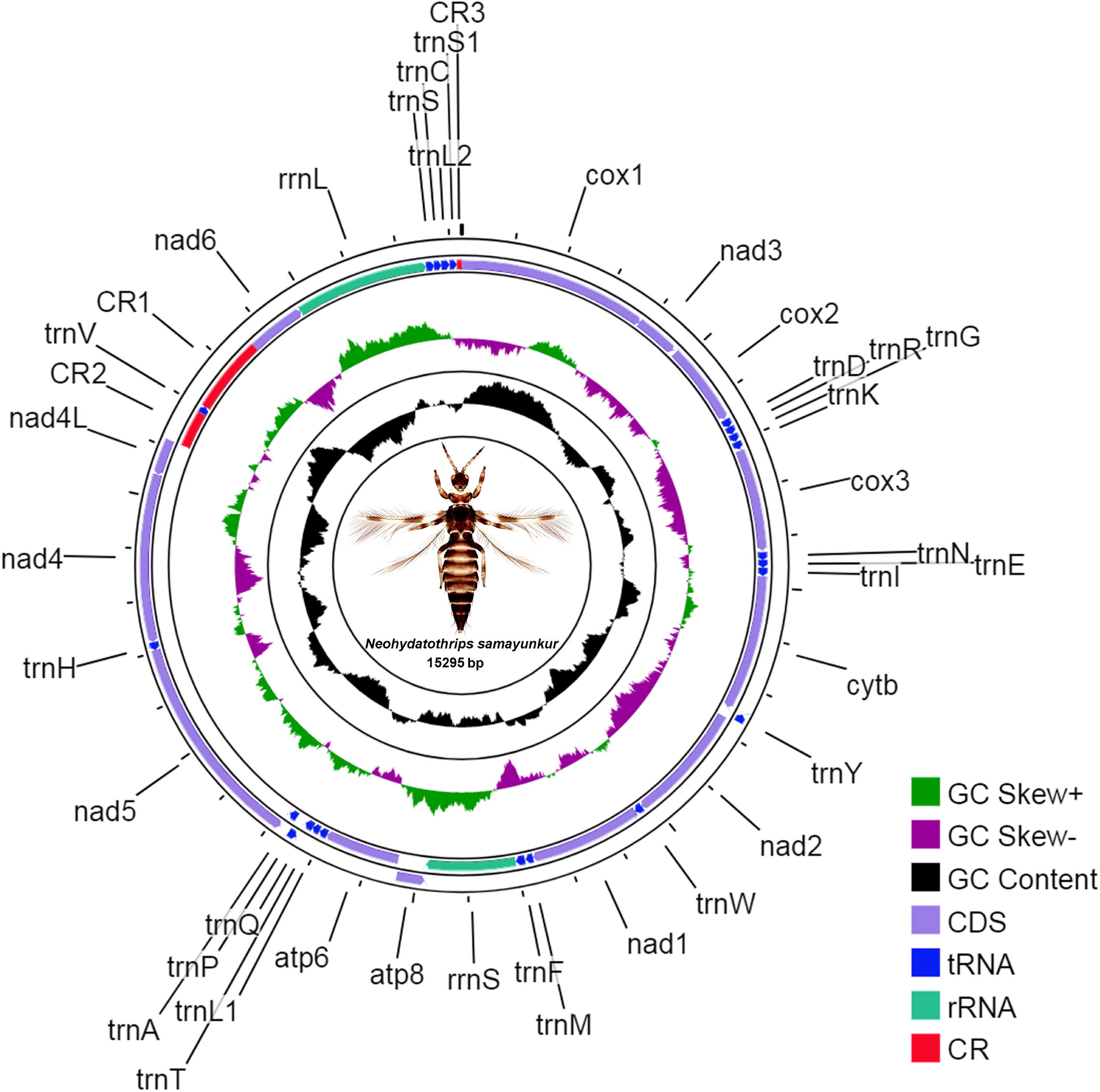
The mitochondrial genome of the marigold thrips, *N. samayunkur.* Direction of gene transcription is indicated by arrows in entire complete genome. PCGs are shown as purple arrows, rRNA genes as sea green arrows, tRNA genes as blue arrows and CR regions as red rectangles. The GC content is plotted using a black sliding window, as the deviation from the average GC content of the entire sequence. GC-skew is plotted using a colored sliding window (green and orchid color), as the deviation from the average GC-skew of the entire sequence. The figure was drawn using CGView online server (http://stothard.afns.ualberta.ca/cgview_server/) with default parameters. The species photograph was taken by second author (KT) using Leica Microscope DM1000 with Leica software application suite (LAS EZ) and edited manually in Adobe Photoshop CS 8.0.

**Table 1.**
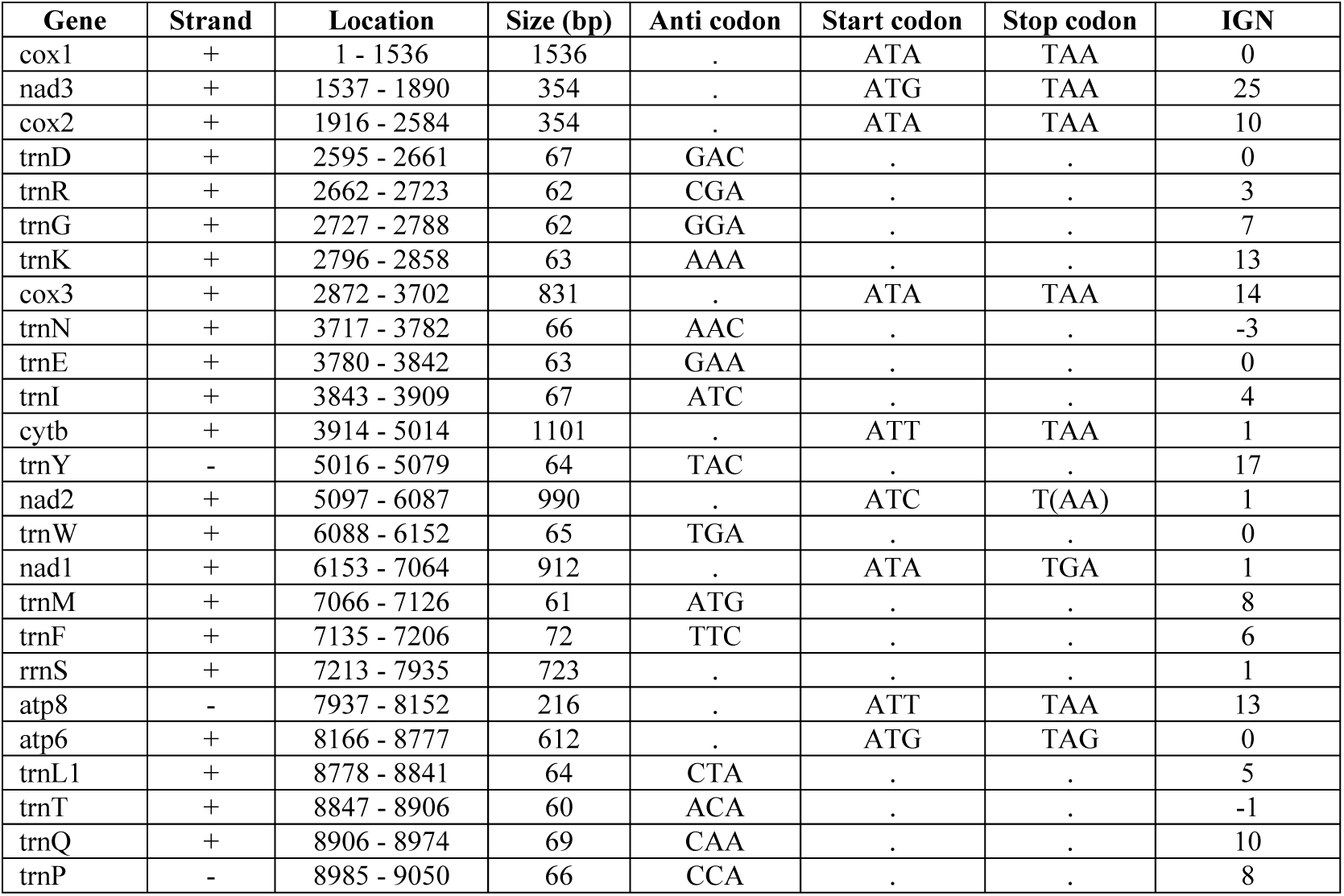
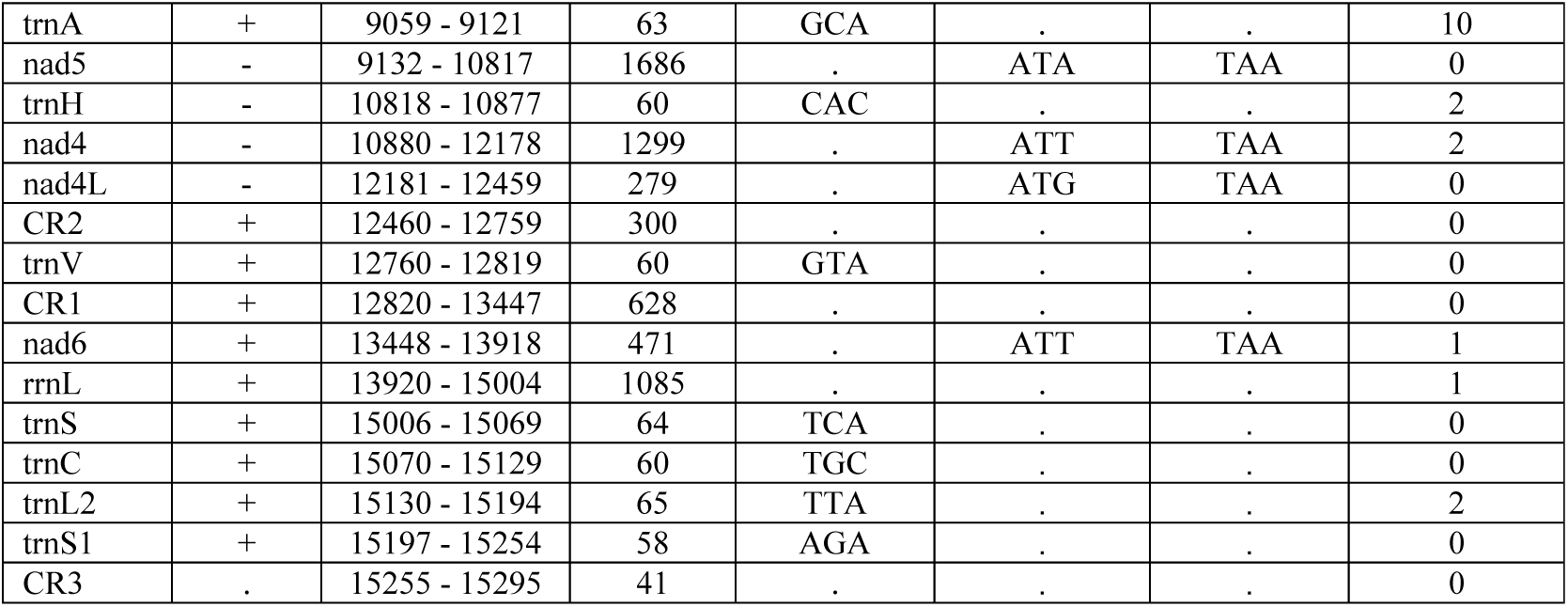
List of annotated mitochondrial genes of *Neohydatothrips samayunkur* and its characteristic features. The protein coding and ribosomal RNA genes are represent by standard nomenclature, tRNAs are represented as trn followed by the IUPAC-IUB single letter amino acid codes. (+) values in strand represent as heavy (H) and (-) values represent as light (L). IGN represents (+) values as intergenic nucleotides and (-) values as overlapping regions. CR represents the control region.

**Table 2.** Composition and skewness in different Thysanoptera mitogenomes considered for comparative analysis.

| Species | Size(bp) | A% | G% | T% | C% | GC% | AT% | ATskewness | GCskewness |
| --- | --- | --- | --- | --- | --- | --- | --- | --- | --- |
| **Whole mtgenome** |  |  |  |  |  |  |  |  |  |
| *N. samayunkur* | 15,295 | 40.25 | 10.98 | 37.17 | 11.60 | 22.58 | 77.42 | 0.04 | -0.03 |
| *A. obscurus* | 14,890 | 38.38 | 11.27 | 39.75 | 10.60 | 21.87 | 78.13 | -0.02 | 0.03 |
| *S. dorsalis* EA1 | 15,343 | 39.12 | 11.61 | 36.62 | 12.64 | 24.26 | 75.74 | 0.03 | -0.04 |
| *S. dorsalis* SA1 | 15,204 | 39.83 | 11.18 | 37.56 | 11.42 | 22.60 | 77.39 | 0.03 | -0.01 |
| *F. intonsa* | 15,215 | 41.24 | 11.06 | 34.68 | 13.01 | 24.07 | 75.93 | 0.09 | -0.08 |
| *F. occidentalis* | 14,889 | 40.98 | 11.35 | 36.62 | 11.06 | 22.41 | 77.59 | 0.06 | 0.01 |
| *T. imaginis* | 15,407 | 43.85 | 10.47 | 32.72 | 12.96 | 23.43 | 76.57 | 0.15 | -0.11 |
| **PCG** |  |  |  |  |  |  |  |  |  |
| *N. samayunkur* | 10,956 | 39.55 | 10.81 | 37.58 | 12.07 | 22.87 | 77.13 | 0.03 | -0.06 |
| *A. obscurus* | 10,814 | 36.85 | 11.60 | 40.23 | 11.32 | 22.91 | 77.09 | -0.04 | 0.01 |
| *S. dorsalis* EA1 | 10,954 | 38.06 | 11.92 | 36.53 | 13.48 | 25.41 | 74.59 | 0.02 | -0.06 |
| *S. dorsalis* SA1 | 10,960 | 38.92 | 11.37 | 37.66 | 12.04 | 23.41 | 76.58 | 0.02 | -0.03 |
| *F. intonsa* | 11,005 | 39.95 | 11.39 | 34.58 | 14.08 | 25.48 | 74.52 | 0.07 | -0.11 |
| *F. occidentalis* | 10,850 | 39.82 | 11.62 | 36.72 | 11.84 | 23.47 | 76.53 | 0.04 | -0.01 |
| *T. imaginis* | 10,899 | 42.76 | 10.16 | 32.86 | 14.23 | 24.39 | 75.61 | 0.13 | -0.17 |
| **tRNA** |  |  |  |  |  |  |  |  |  |
| *N. samayunkur* | 1,397 | 43.16 | 9.88 | 37.65 | 9.31 | 19.18 | 80.82 | 0.07 | 0.03 |
| *A. obscurus* | 1,412 | 39.87 | 10.69 | 39.80 | 9.63 | 20.33 | 79.67 | 0.00 | 0.05 |
| *S. dorsalis* EA1 | 1,418 | 40.69 | 10.93 | 37.52 | 10.86 | 21.79 | 78.21 | 0.04 | 0.00 |
| *S. dorsalis* SA1 | 1,419 | 41.51 | 10.43 | 38.13 | 9.94 | 20.37 | 79.63 | 0.04 | 0.02 |
| *F. intonsa* | 1,386 | 43.51 | 10.68 | 35.79 | 10.03 | 20.71 | 79.29 | 0.10 | 0.03 |
| *F. occidentalis* | 1,376 | 42.44 | 10.47 | 37.43 | 9.67 | 20.13 | 79.87 | 0.06 | 0.04 |
| *T. imaginis* | 1,457 | 43.72 | 9.54 | 36.72 | 10.02 | 19.56 | 80.44 | 0.09 | -0.02 |
| **rRNA** |  |  |  |  |  |  |  |  |  |
| *N. samayunkur* | 1,808 | 44.97 | 11.73 | 34.35 | 8.96 | 20.69 | 79.31 | 0.13 | 0.13 |
| *A. obscurus* | 1,812 | 43.16 | 11.70 | 36.59 | 8.55 | 20.25 | 79.75 | 0.08 | 0.16 |
| *S. dorsalis* EA1 | 1,775 | 43.21 | 11.89 | 34.99 | 9.92 | 21.80 | 78.20 | 0.11 | 0.09 |
| *S. dorsalis* SA1 | 1,777 | 45.36 | 11.65 | 34.44 | 8.55 | 20.20 | 79.80 | 0.14 | 0.15 |
| *F. intonsa* | 1,699 | 47.15 | 11.30 | 32.02 | 9.54 | 20.84 | 79.16 | 0.19 | 0.08 |
| *F. occidentalis* | 1,848 | 45.94 | 12.18 | 33.93 | 7.95 | 20.13 | 79.87 | 0.15 | 0.21 |
| *T. imaginis* | 1,876 | 47.65 | 10.77 | 32.14 | 9.43 | 20.20 | 79.80 | 0.19 | 0.07 |
| **Control Region** |  |  |  |  |  |  |  |  |  |
| *N. samayunkur* | 969 | 34.57 | 14.34 | 37.15 | 13.93 | 28.28 | 71.72 | -0.04 | 0.01 |
| *A. obscurus* | 145 | 25.52 | 8.97 | 62.76 | 2.76 | 11.72 | 88.28 | -0.42 | 0.53 |
| *S. dorsalis* EA1 | 820 | 40.12 | 9.51 | 39.02 | 11.34 | 20.85 | 79.15 | 0.01 | -0.09 |
| *S. dorsalis* SA1 | 767 | 35.33 | 9.26 | 43.55 | 11.86 | 21.12 | 78.88 | -0.10 | -0.12 |
| *F. intonsa* | 942 | 41.72 | 7.86 | 38.22 | 12.21 | 20.06 | 79.94 | 0.04 | -0.22 |
| *F. occidentalis* | 595 | 40.34 | 7.90 | 43.70 | 8.07 | 15.97 | 84.03 | -0.04 | -0.01 |
| *T. imaginis* | 900 | 47.56 | 16.67 | 25.22 | 10.56 | 27.22 | 72.78 | 0.31 | 0.22 |

### Protein-coding genes and Relative Synonymous Codon Usage (RSCU)

The mitochondrial genome of *N. samayunkur* contained 13 PCGs, starting with ATN (five with ATA, four with ATT, three with ATG and one with ATC) initiation codons. The stop codon TAA is detected for 10 PCGs, TGA for *nad1* and TAG for *atp6*, while an incomplete termination stop codon is present at *nad2.* The TGA stop codon in *nad1* was only observed in *N. samayunkur*. The *atp6* gene was terminated with TAG stop codon in *N. samayunkur* while TAA was observed in other thrips species. This type of incomplete termination codon was also observed in *nad2* (*T. imaginis*), *atp8* (*A. obscurus* and *T. imaginis*) and *nad4* (*S. dorsalis* EA1 and SA1, *F. occidentalis*, *F. intonsa* and *T. imaginis*) (Table 1). The presence of an incomplete stop codon is well documented in metazoan mitogenomes and which are supposed to be completed via post-transcriptional polyadenylation^20^ The comparative analysis of nucleotide composition in thrips revealed that the average adenine (A) composition was 33.96% within all studied PCGs and resulted highest (56.15%) in *nad4L* gene of *A. obscurus* and lowest (22.23%) in *nad4* gene of *T. imaginis.* The analysis resulted highest adenine composition ranging from 34.13% to 45.78% in most of the PCGs in *T. imaginis* as compared to other thrips species. Further the adenine composition was much higher in three PCGs (*nad5*, *nad4* and *nad4L*) in *N. samayunkur* than the average. The average thiamine (T) composition in all PCGs of thrips mitogenomes was 42.93% and resulted highest in *A. obscurus*, (41.32% to 50.58%) for most of the PCGs (*cox1*, *nad3*, *cox3*, *cytb*, *nad2*, *nad1*, *atp6* and *nad6*). The average guanine (G) composition was 10.87% in all PCGs of thrips mitogenomes and *S. dorsalis* east Asian (EA1) strain was observed with highest, (9% to 12.88%) in five of the PCGs (*nad3*, *cytb*, *nad2*, *nad1*, and *nad4L*). The average cytosine (C) composition was 12.24% for thrips mitogenomes PCGs and highest, (12.06% to 15.99%) in five PCGs (*cox2*, *cytb*, *nad2*, *nad1*, and *nad6*) of *T. imaginis.* The average GC composition within all PCGs was 23.11% in the studied thrips mitogenomes. The GC composition was highest in seven PCGs (*nad3*, *cox2*, *cytb*, *nad1*, *atp6*, *nad4L*, and *nad6*) in *S. dorsalis* EA1 ranging from 21.28% to 28.46% (Table S2, Fig. 2). The comparative codon usage analysis of *N. samayunkur* revealed that Leucine (L), Alanine (A), Arginine (R), Glycine (G), Proline (P), Threonine (T) and Valine (V) were the most frequently utilized amino acids. The Relative Synonymous Codon Usage (RSCU) analysis of PCGs revealed that the codon distributions of six thrips species were consistent, and each amino acid has almost equal contents in different species (Fig. 3). Sequence saturation analysis of PCGs in *N. samayunkur* mitogenome showed the increase of frequency of both transitions and transversions linearly along with the divergence value (Table S4, Fig. 4).

**Figure 2.**
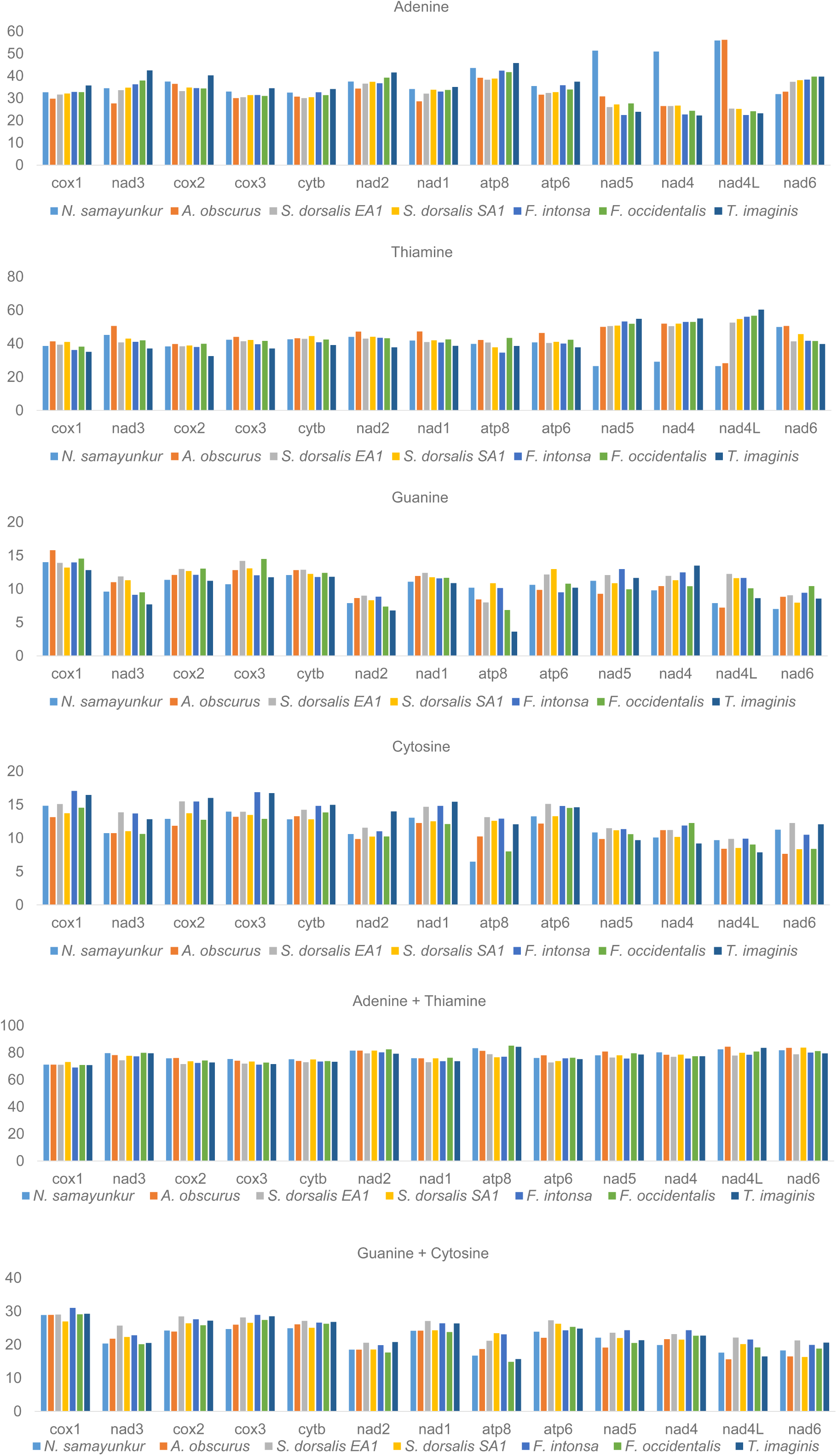
Comparative analysis of nucleotide composition of PCGs within the studied thrips species. The figure was drawn using MEGA6, and edited manually in Microsoft Excel.

**Figure 3.**
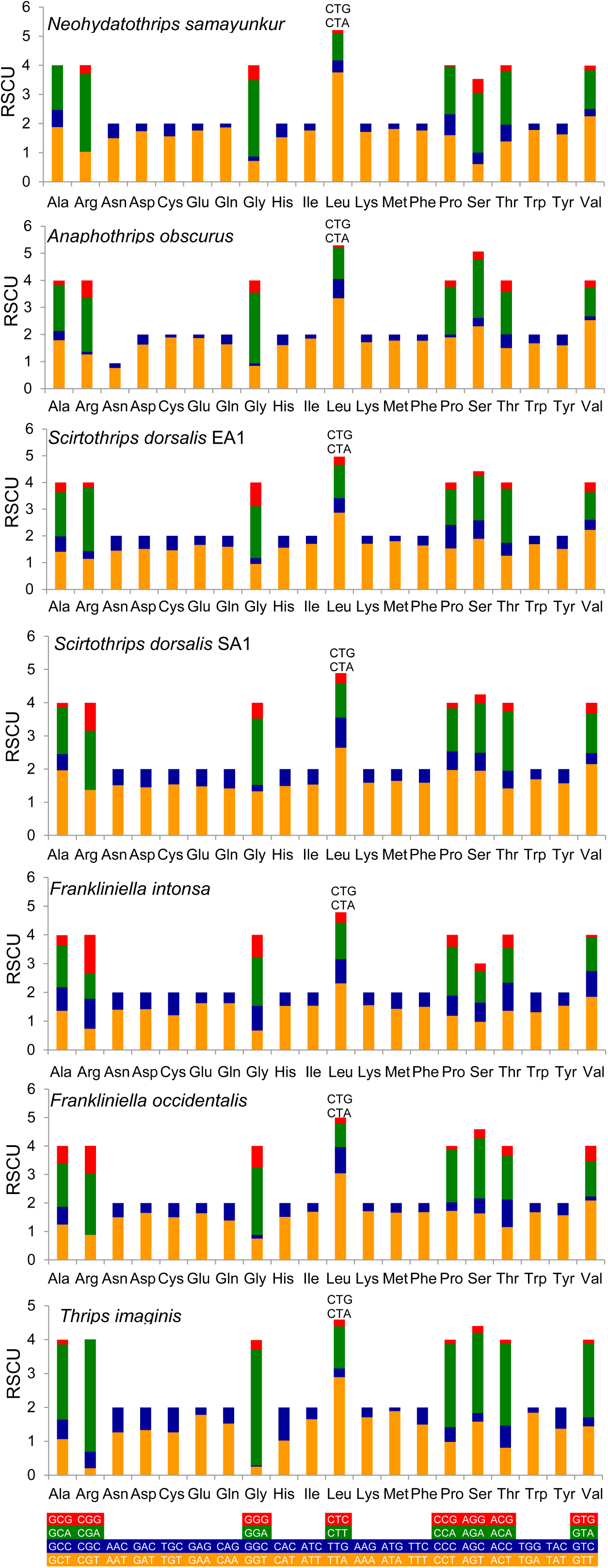
The Relative Synonymous Codon Usage (RSCU) of the mitochondrial genome of *N. samayunkur* and other studied thrips species. Codons indicated above the Leucine (Leu) bar are represent same codon present in all studied thrips mitogenomes. The figure was drawn using MEGA6, and edited manually in Microsoft Excel.

**Figure 4.**
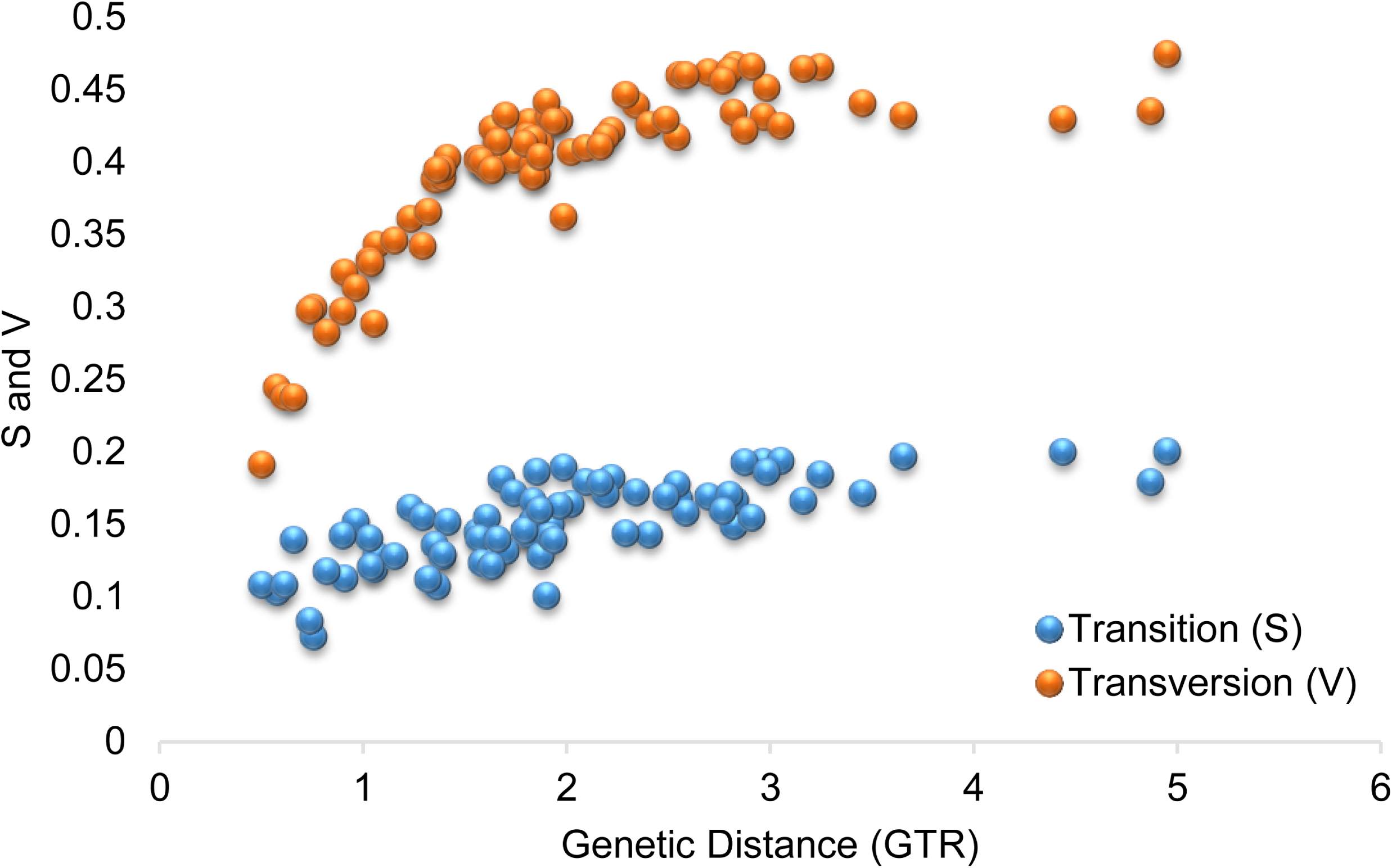
Transition (S) and transversion (V) saturation plots for dataset (protein-coding genes) of *N. samayunkur*. The figure was drawn using DAMBE5 software with default parameters.

### Ribosomal and transfer RNA genes

The mitogenome of *N. samayunkur* comprises two rRNA genes as in other insects mitogenomes. The large ribosomal gene (16S rRNA) was 1085 bp long, and located between *nad6* and tRNA Serine (S). The small ribosomal gene (12S rRNA) was 723 bp long, and located between tRNA Phenyalanine (F) and *atp8* gene (Table 2). The AT content (79.31%) of two rRNAs were observed within the range from 78.20% (*S. dorsalis* EA1) to 79.87% (*F. occidentalis*) of other thrips mitogenomes. Both AT skewness (0.13) and GC skewness (0.13) were positive, that is similar to other previously sequenced thrips mitogenomes.

The *N. samayunkur* mitogenomes contained complete set of 22 tRNAs (ranging from 58 to 72 nucleotides in length) with a total length of 1,397 bp. This region was highest in AT richness among thrips mitogenomes, represented by 80.82%, with positive AT skewness (0.07), and GC skewness (0.03) (Table 2). Further, 19 tRNA genes were coded by the H-strand and rest three (*trnY*, *trnP and trnH*) by the L-strand. The *trnY* and *trnF* were coded by L-strand and H-strand respectively in all the thrips species except *S. dorsalis* SA1. The *trnS* was coded by H-strand in all thrips species except *T. imaginis.* The *trnP* was coded by H-strand in *A. obscurus* and *S. dorsalis* while coded by L-strand in other thrips species. All tRNA showed the typical cloverleaf secondary structure except *trnV and trnS* without DHU stem and loop (Fig. 5). The absence of DHU stem and loop in *trnV* was consistent in all the thrips species mitogenomes. The mitogenomes of *N. samayunkur* along with five thrips species varied considerably in the arrangement of tRNA genes. The six of the 22 tRNAs genes (*trnG*, *trnK*, *trnY*, *trnW*, *trnF*, and *trnH*) were found to be conserved, among the five thrips species in their locations relative to a protein-coding or rRNA gene upstream or downstream (Fig. 6A). The four tRNA genes (*trnL2*, *trnA*, *trnC*, and *trnV*) were found to be variable in *N. samayunkur* contrary to their conserved location in other thrips species. The *trnL2* was located upstream of *cox2* in other thrips species, whereas, it was translocated between *trnC* and *trnS2* in *N. samayunkur.* The *trnA*-*F* were located upstream 12S *rRNA*, however, *trnA* was translocated upstream of *nad5* gene in *N. samayunkur.* The *trnC* was upstream of *nad6* gene and translocated to between *trnS1 and trnL2* in *N. samyunkur.* The *trnV* was upstream of 16S *rRNA*, however, it was translocated between two control regions (CR1 and CR2) in *N. samayunkur* (Fig. 6A).

**Figure 5.**
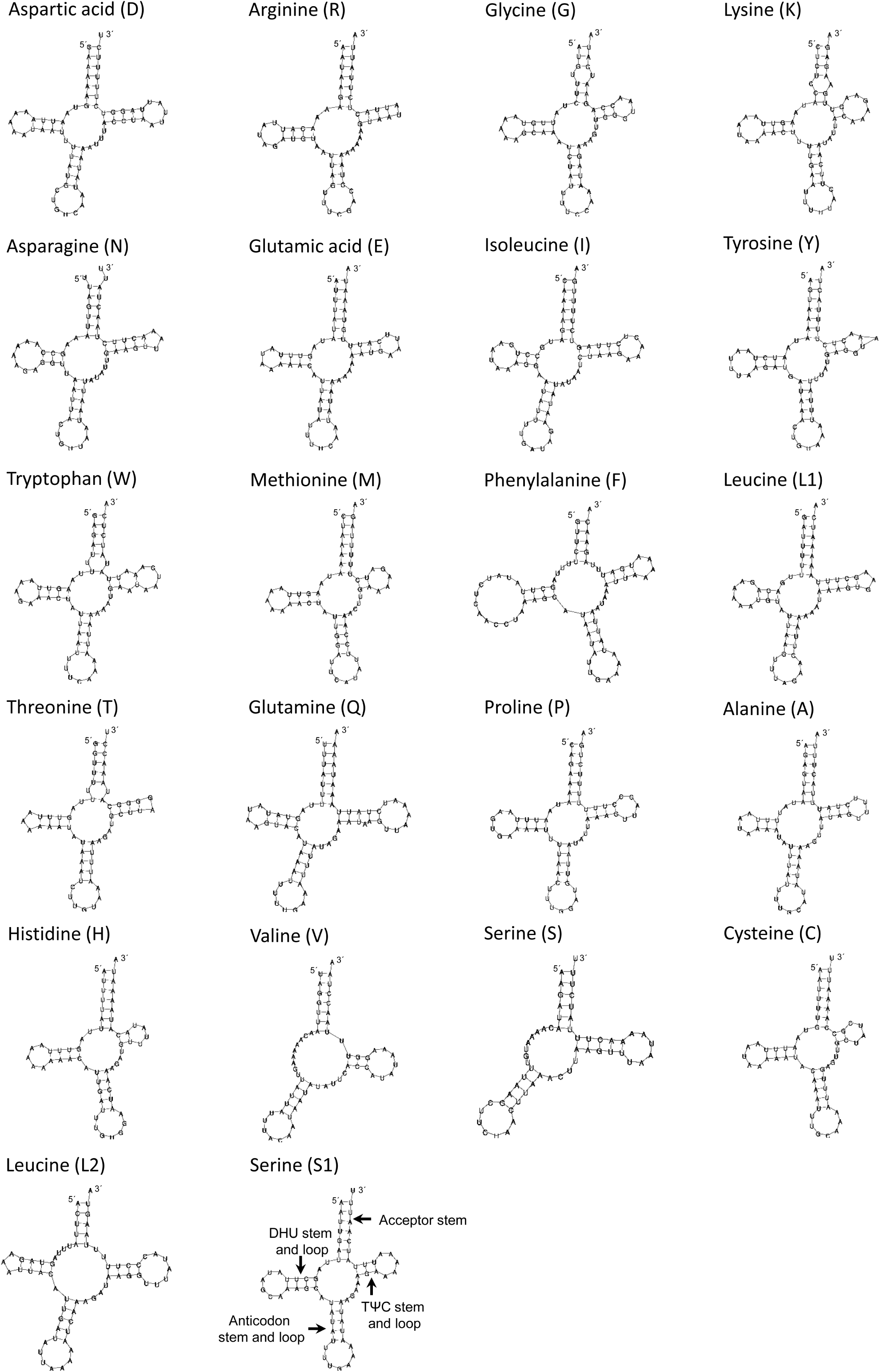
Putative secondary structures of the 22 tRNA genes of the *N. samayunkur* mitogenome. The tRNAs are represented by full names and IUPAC-IUB single letter amino acid codes. The details of stem and loop is mentioned for one tRNA Serine (S1) which is applicable for all tRNAs secondary structures. The secondary structure of tRNAs were predicted by MITOS online server (http://mitos.bioinf.uni-leipzig.de/index.py) and edited manually in Adobe Photoshop CS 8.0.

**Figure 6.**
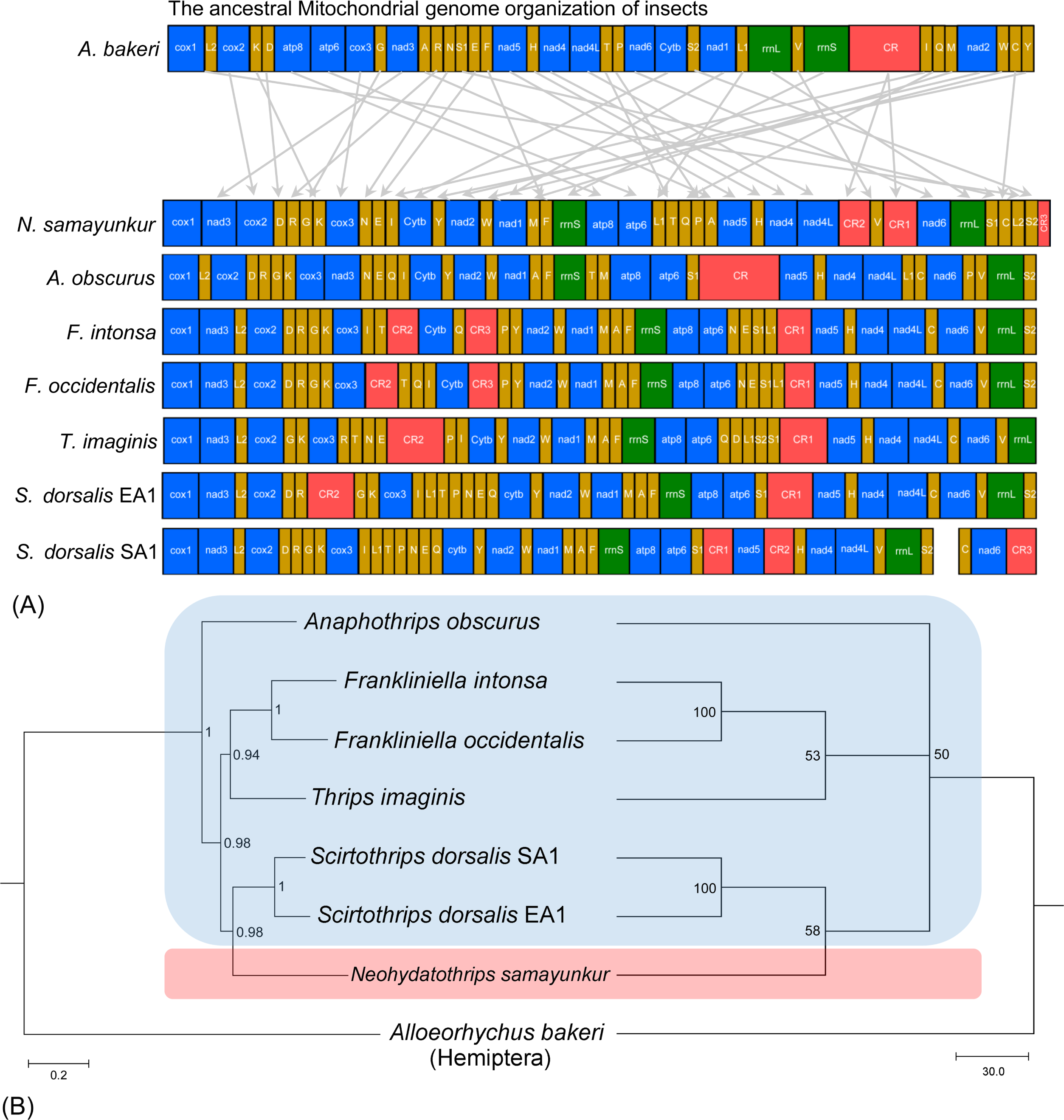
(A) Mitogenome arrangement of *N. samayunkur* with respect to other thrips species. Ancestor here denotes *A. bakeri* (Hemiptera). The protein coding and ribosomal RNA genes are represent by standard nomenclature with blue and green color respectively. The tRNAs are represented by the IUPAC-IUB single letter amino acid codes and shown in golden color. The control regions are represented by CR with red color. Grey arrows indicates the gene rearrangement between the *N. samayunkur* with ancestor species. The figure was generated manually in Microsoft Excel and edited in Adobe Photoshop CS 8.0. (B) Phylogenetic tree (left side) inferred from nucleotide sequences of 13 PCGs using Bayesian Inference method in MrBayes v3.2. The tree is drawn to scale with bayesian posterior probability values indicated along with the branches. Phylogenetic tree (right side) inferred from nucleotide sequences of 13 PCGs using maximum likelihood method in PAUP (1000 bootstrap replicates). The tree is drawn to scale with bootstrap values indicated along with the branches. The bluish color represent the species clustering of subfamily Thripinae and reddish color represent the species of subfamily Sericothripinae. The figure was edited in Adobe Photoshop CS 8.0.

### Overlapping and intergenic spacer regions

The mitogenome of *N. samayunkur* contained two overlapping regions with a total length of 4 bp. The first overlapping region with three nucleotides was observed between tRNA Asparagine (N) and tRNA Glutamic Acid (E). The second overlapping region with one nucleotide was depicted between tRNA Threonine (T) and tRNA Glutamine (Q). The mitogenome of *N. samayunkur* had 24 intergenic spacer regions in a total of 165 bp, varying from 1 to 25 bp in length. There are eight major intergenic spacers of at least 10 base pair in length (Table 1). The longest intergenic spacer (25bp) is observed between the *nad3* and *cox2* gene, with an extremely high AT content. The AT richness in intergenic spacers had also been observed in other thrips mitogenomes. The comparative analysis of *N. samayunkur* with other thrips mitogenomes revealed that *S. dorsalis* SA1 had the highest overlapping region (15) with a total length of 86 bp, varying from 1 to 49 bp and lowest intergenic spacer (13) with a total of 342 bp, varying from 1 to 85 bp (Table S3).

### Control Regions (CRs)

The *N. samayunkur* mitogenome contained three control regions (CR1, CR2, and CR3). The CR1 (628 bp) was lied between *trnV* and *nad6*, CR2 (300 bp) located between the *nad4L* and *trnV*, and CR3 (41 bp) located between *trns1* and *cox1.* Further, the CR1 had 89.33% and 63.41% sequence similarity with CR2 and CR3 respectively, indicating a possible duplication and translocation of control region. The total length of CR is 969 bp which is the highest in comparison with the existing mitogenomes of thrips. It is to note that the number and locations of control regions (CRs) or A+T rich regions varied among different thrips species mitogenomes. *A. obscurus* had one CR, *T. imaginis* and *S. dorsalis* EA1 had two CRs, whereas the two *Frankliniella* species (*F. intonsa* and *F. occidentalis*) and *S. dorsalis* SA1 had three CRs respectively. The location of CR1 upstream of *nad5* gene was suggested to be ancestral condition of the thrips species in subfamily Thripinae^19^. However, the location of CRs in *N. samayunkur* is not in concordance with other thrips mitogenomes in subfamily Thripinae. The arrangement of CRs in thrips taxa seems to be highly variable and need more mitogenomes data from different taxa to clarify the evolution of CR in thrips.

### Phylogenetic analyses

Both the Maximum likelihood (ML) and Bayesian Inference (BI) phylogenetic trees generated resulted similar topologies (Fig. 6B). The two *Frankliniella* species along with *T. imaginis* were clustered together. The phylogenetic analyses showed that *N. samayunkur* was clustered with *S. dorsalis* (SA1 and EA1) which is in coherence with the results of multi-gene molecular data^8,9^ and morphological data^10^. The suborder Terebrentia is classified into eight families with four subfamilies^21^ which is most excepted views in taxonomic fraternity. However, an alternative view proposed 28 families in suborder Terebrentia based on highly conserved taxonomic characters and elevated the subfamily Sericothripinae to family rank^22^. This alternative proposal was refuted by the molecular data^8,23^. Till date, molecular phylogenetic studies on thrips is in its early stages due to lack of large scale data with more taxonomic sampling. Thus, the generation of comprehensive molecular data on families/subfamilies need to be investigated. The present mitogenomic analysis indicated that the *S. dorsalis* (subfamily Thripinae) was close to *N. samayunkur* (subfamily Sericothripinae) than to other Thripinae taxa. The close relationships between *Scirtothrips* genus-group and Sericothripinae was also supported by the following shared taxonomic characters^10^: presence of closely spaced rows of microtrichia on lateral thirds of abdominal tergites; median pair of tergal setae close together; campaniform sensilla absent on tergite IX, tergite X not split longitudinally. We concluded that more molecular data on the diverse thrips species from different hierarchical level is needed, to understand the phylogenetic and evolutionary relationships among them.

## Materials and Methods

### Sample collection and DNA extraction

The adult specimens of *N. samayunkur* were collected from the Odisha State of India. Specimens were morphologically identified by the second author with help on available taxonomic keys^2^, and preserved in absolute ethyl alcohol at –30°C in Centre for DNA Taxonomy, Molecular Systematics Division, Zoological Survey of India, Kolkata. The genomic DNA was extracted from using the DNeasy DNA Extraction kit (QIAGEN) following the manufacturer’s standard protocol. The concentration of DNA was analysed on a Qubit fluorometer using a dsDNA high-sensitivity kit (Invitrogen). Further, we checked the quality of the extracted DNA by on 0.8% agarose gel electrophoresis.

### Mitogenome sequencing and assembly

The whole genome library of genomic DNA was sequenced using the Illumina Hiseq2500 (2 × 150 base paired-end reads) (Illumina, USA) platform which yielded ~23 million reads. The paired-end library was constructed according to the standard protocols using the TruSeq DNA Library Preparation kit. Raw sequencing reads were trimmed and filtered according to quality using NGS-Toolkit by removing adapter contamination and low-quality reads with base N’s or more than 70% of bases with a quality score <20. The obtained high quality reads were screened out using Burrows-Wheeler Alignment (BWA) tool^24^ and then assembled with SPAdes 3.9.0^25^, using default parameters considering *S. dorsalis* mitochondrial genome (NC_025241.1) as trusted contig. The high quality ~19 million reads was screened out using BWA aligner with default parameters. Out of 19 million reads, 0.88% (~1.7 million) of the reads got aligned to the reference used. The aligned reads were considered for denovo mitochondrial genome assembly.

### Genome annotation, visualization, and comparative analysis

The assembled mitogenome was annotated using widely used MITOS web-server (http://mitos.bioinf.uni-leipzig.de/index.py) to estimate the location of protein coding regions, tRNAs, rRNAs and their secondary structures^26^. The FASTA-formatted mitochondrial genome assembly was performed by aligning the contigs against the non-redundant nucleotide database of NCBI (National Center for Biotechnology Information) using the BLASTn search algorithm (http://blast.ncbi.nlm.nih.gov/Blast). The location of PCGs and rRNAs was confirmed manually by BLASTn, BLASTp and ORF Finder in NCBI (https://www.ncbi.nlm.nih.gov/orffinder/). The nucleotide sequences of the Protein Coding Genes (PCGs) were initially translated into putative proteins on the basis of the invertebrate mitochondrial DNA genetic code. The initiation and termination codons were identified in ClustalX using other Thrips reference sequences^27^. MEGA6 was used for alignment of the homologous sequences of *N. samayunkur* with the other thrips species^28^. The complete annotated mitogenome was submitted to NCBI GenBank using Sequin tool (http://www.ncbi.nlm.nih.gov/Sequin/) to acquire the unique accession number. The circular map of *N. samayunkur* mitogenome was illustratred by CGView online server (http://stothard.afns.ualberta.ca/cgview_server/) with default parameters^29^. We analysed in the complete mitogenome of *N. samayunkur* and other available thrips mitogenomes in GenBank database (Table S1) to estimate the nucleotide composition, AT and GC skewness. Further, the nucleotide composition of PCGs in *N. samayunkur* and other available thrips mitogenomes were compared. MEGA6 was used for estimation of nucleotide composition, codon usages, relative synonymous codon usage (RSCU) and composition of skewness with the following formula: AT skew = (A – T)/(A + T) and GC skew = (G – C)/(G + C)^30^. The DAMBE5 software was used to test the sequence substitution saturation of PCGs in *N. samayunkur* mitogenome^31^. The typical cloverleaf secondary structure of transfer RNA (tRNA) genes were predicted by MITOS were further confirmed using the tRNAscan-SE (http://lowelab.ucsc.edu/tRNAscan-SE/)^32^ and ARWEN 1.2^33^. The Overlapping sequence and intergenic spacer regions of *N. samayunkur* and other published thrips mitogenomes were compared in terms of length and location. Further, the homology of control region with other thrips mitogenomes were determined through sequence alignment using Clustal Omega^27^.

### Phylogenetic analyses

We retrieved the six completed mitogenomes of thrips species available in GenBank for the phylogenetic inference. *Alloeorhychus bakeri* (Hemiptera) was used as an out-group^34^. To reconstruct the phylogenetic relationship among thrips species, the nucleotide sequences of the 13 PCGs were aligned and concatenated to form dataset. Each PCG was aligned individually using the MAFFT algorithm in the TranslatorX online platform under the L-INS-i strategy based on codon-based multiple alignments^35^. The poorly aligned sites were removed from the protein alignment using GBlocks within the TranslatorX with default settings. The dataset of 13 PCGs (11,148 bp residues) was concatenated using SequenceMatrix v1.7.845^36^ The optimal substitution models for each dataset were selected by PAUP under the automated model selection criteria^37^. The dataset was analyzed using maximum likelihood (ML) method implemented in PAUP, and Bayesian inference (BI) method implemented in MrBayes 3.2^38^. For ML analyses, the bootstrap analysis of 1,000 replicates was performed in PAUP with GTR+G+I model with following settings (LSet nst = 6; rclass = abcdef; rmatrix = 1.05493, 7.3156661, 3.2038249, 7.6517349, 9.8362651; basefreq = 0.33386688, 0.12022167, 0.10899419; rates = gamma; shape = 1.2214965; pinv = 0.27577905). For BI analyses, two simultaneous runs of 10 million generations were conducted for the dataset using GTR+G+I model and trees were sampled every 1,000 generations, with the first 25% discarded as burn-in. Stationary was considered to be reached when the average standard deviation of split frequencies was below 0.01. The phylogenetic tree was visualized and edited using FigTree v1.4.2 (http://tree.bio.ed.ac.uk/software/figtree/)^39^.

## Acknowledgement

The authors are thankful to the Director, Zoological Survey of India, Kolkata, for providing necessary facilities, constant support and encouragement throughout the study. The study is financially supported by Zoological Survey of India, Kolkata, Ministry of Environment Forest and Climate Change under National Faunal Genome Resources (NFGR) Program. This work is a part of the Ph. D thesis of the RC.

## Authors’ contribution

K.T., V.K. and D.S. collected specimens, K.T. and V.K. conceived and designed the experiment, K.T. performed taxonomic identification of the thrips species and captured photographs, V. K. and K. C. contributed chemicals, K.T., S. K., R.C. generated DNA data, V. K., K.T., and S. K., analysed the data, wrote the manuscript text, and prepared the figures, all authors reviewed the manuscript.

## Additional information

Supplementary information accompanies this paper at

